# Orphan Non-Coding RNAs Drive Tumorigenesis and Enable Pre-Diagnostic Detection in HPV-Negative Head and Neck Cancer

**DOI:** 10.64898/2025.12.20.695703

**Authors:** Hoang C. B. Nguyen, Lishi Li, Mehran Karimzadeh, Jack Ghanam, Jeffrey Wang, Yana Al Inaya, Gjystina Lumaj, Fereydoun Hormozdiari, Babak Alipanahi, Allen L. Feng, Jeremy D. Richmon, Daniel G. Deschler, Mark A. Varvares, Daniel L. Faden, Derrick T. Lin, Hani Goodarzi

**Author notes:** Co-corresponding authors, Hoang C. B. Nguyen, MD PhD, Department of Otolaryngology, Head and Neck Surgery, Harvard Medical School, Massachusetts Eye and Ear, 243 Charles Street, Boston, MA 02114, Derrick T. Lin, MD, FACS, Daniel Miller Professor and Associate Chair, Department of Otolaryngology, Head and Neck Surgery, Harvard Medical School, Massachusetts Eye and Ear, Massachusetts General Hospital, 243 Charles Street, Boston, MA 02114, Hani Goodarzi, PhD, Core Investigator, Arc Institute, Associate Professor, Department of Biochemistry & Biophysics, University of California, San Francisco.

## Abstract

HPV-negative head and neck squamous cell carcinoma (HNSCC) lacks reliable biomarkers for early detection and molecular stratification. We recently described orphan non-coding RNAs (oncRNAs) as a class of cancer-emergent small RNAs that arise from cryptic promoters largely silent in normal tissues. Here, we define the comprehensive landscape of oncRNAs in HNSCC. Using TCGA and CPTAC-3 data, we demonstrate that these transcripts are not only tumor-specific but provide digital fingerprints that accurately stratify tumors by anatomical site. Moving from association to function, we investigated whether these emergent transcripts actively drive tumorigenesis. We focused on oncRNA64585, a transcript highly enriched in oral cavity tumors and associated with tumor progression. Its depletion significantly impaired proliferation in vitro and suppressed tumor growth in syngeneic murine models. Mechanistically, transcriptomic and chromatin accessibility profiling revealed that oncRNA64585 regulates transcriptional programs governing cell cycle progression and apoptosis through interaction with the Polycomb Repressive Complex 2 (PRC2). Finally, we demonstrate the clinical utility of these signatures: oncRNA profiles in plasma predicted oral cavity cancer with >90% accuracy up to four years prior to clinical diagnosis. These findings establish oncRNAs as active drivers of HNSCC biology and potent tools for early detection and therapeutic targeting.

## INTRODUCTION

Head and neck squamous cell carcinoma (HNSCC) is a clinically and molecularly heterogeneous malignancy ^1–3^. Despite recent therapeutic advances, patients with HPV-negative disease continue to suffer from recurrent disease following curative treatment attempts ^3–6^. These patients often present with locally advanced disease, afflicted with treatment-related morbidity, and lack biomarkers for early diagnosis or treatment stratification ^7–11^. Thus, there is a need for new molecular tools that can both improve early detection and inform novel therapeutic strategies.

Recent studies have begun to reveal the critical roles of cancer-emergent non-coding RNAs in cancer progression, but much of the non-coding transcriptome remains unexplored, especially in the context of head and neck cancers. We recently described a novel class of tumor-specific small non-coding RNAs—termed orphan non-coding RNAs (oncRNAs)—which are detectable in tumor tissues but largely absent in normal tissues and blood^12,13^. These newly annotated RNAs arise from activation of cryptic promoters following epigenomic reprogramming in cancer or mis-processing of longer precursor RNAs into smaller but stable non-coding RNAs. We have previously shown that cancer cells can tap into the pool of these cancer-emergent molecules to engineer new oncogenic and pro-metastatic pathways^12,14^. These cancer-emergent programs represent previously unrecognized drivers of tumor progression, and their absence in normal tissues highlights them as attractive therapeutic targets with minimal risk of on-target toxicity.

In addition to their role as potential drivers of human disease, oncRNAs also provide a unique opportunity for developing non-invasive cancer detection assays. A sizable fraction of oncRNAs are actively secreted from cancer cells ^13^, and given their cancer specificity, their presence in blood points to the presence of underlying cancer cells that are secreting them. We have previously leveraged oncRNA fingerprinting enabled by whole-cell free small RNA profiling of serum or plasma for detection of cancer both in case-control studies^15^ and minimal residual disease settings^16^.

Here, we describe the first comprehensive analysis of this novel class of small non-coding RNAs in HNSCC. We show that oncRNAs stratify tumors by anatomical site, correlate with disease progression, and are detectable in patients’ plasma years prior to cancer diagnosis. Moving from biomarker discovery to functional characterization, we demonstrate that oncRNA64585—a highly enriched transcript in oral cavity tumors—acts as a cancer-emergent neo-regulator of chromatin accessibility, driving tumor proliferation through the modulation of cell cycle and apoptosis programs. Our results establish oncRNAs as versatile molecular tools for the early detection, stratification, and therapeutic targeting of HPV-negative HNSCC.

## RESULTS

### Discovery and Molecular Stratification of HNSCC via Site-Specific oncRNA Fingerprints

We recently described the discovery of oncRNAs as a class of small RNAs that were largely absent in normal tissues but expressed in human cancers based on small RNA sequencing data from the Cancer Genome Atlas (TCGA) and cell-free small RNA data from exoRNA atlas was the basis of this discovery. Additionally, we corroborated the expression of oncRNAs in patient-derived xenografts and cell line models. OncRNAs are not only cancer specific but also present cancer type and sub-type specific patterns of expression, providing robust digital molecular fingerprints of cancer cell identity ^1,16^. In this study, we focused on the discovery, annotation, and characterization of cancer-emergent oncRNAs expressed in HNSCC (**Figure 1A**). Approximately 80,000 species of oncRNAs were uniquely identified from over 500 cancer tissue samples from patients with head and neck cancers compared to adjacent normal tissues in the TCGA dataset (**Figure 1B**). The same oncRNAs were also similarly enriched in an independent cohort of patients from the Clinical Proteomic Tumor Analysis Consortium (CPTAC-3) (**Figure 1C**) ^17^.

**Figure 1.**
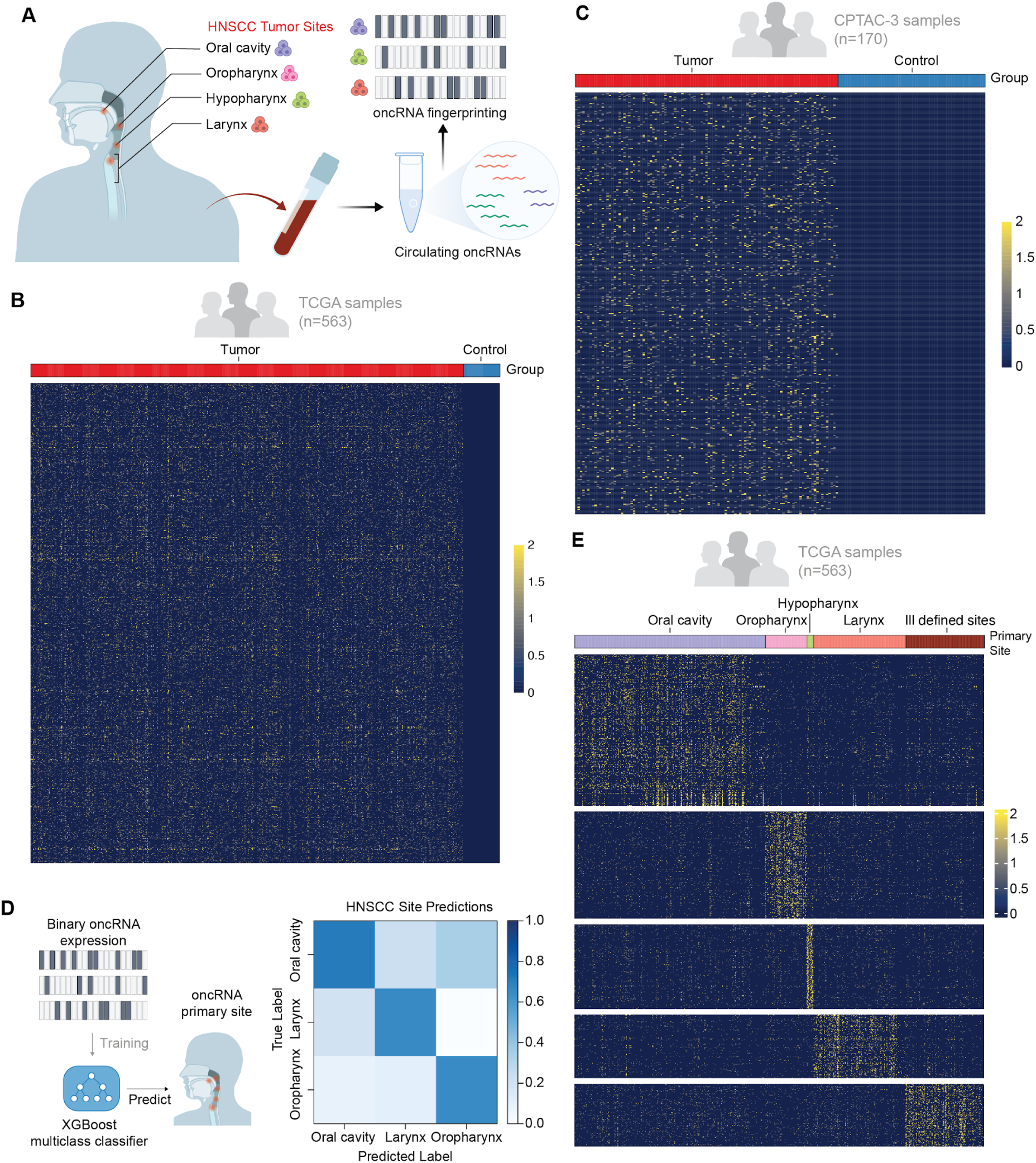
HNSCC can be stratified by their oncRNA profile. **(A)** Schematic of the overarching goal of this study to detect and stratify HNSCC utilizing oncRNA. **(B)** Heatmap showing 83,173 oncRNAs unique to HNSCC from 563 TCGA samples. **(C)** Heatmap showing expression levels of oncRNAs identified from TCGA datasets in 170 CPTAC-3 samples. **(D)** Schematic of XGBoost multiclass classifier to predict HNSCC primary site based on oncRNA profile and the confusion matrix for site-specific classification based on oncRNA presence and absence in each sample. The matrix was row-normalized. **(E)** Heatmap showing oncRNAs specific to different primary sites of HNSCC, including oral cavity, oropharynx, hypopharynx, and larynx from the TCGA dataset.

Since tumor subsites strongly influence prognosis and treatment for patients with head and neck cancers, we next examined whether HNSCC can be stratified by primary tumor site based on the expression of oncRNAs. Even though there is significant overlap in terms of oncRNA expression profile among HNSCC samples, distinct subsets of oncRNAs appear unique to the various sites of origin (**Figure S1A**). We asked whether machine learning models can unbiasedly stratify HNSCC-associated oncRNAs by their tumor origin. We evaluated the performance of an XGBoost multiclass classifier trained on the binary detection of oncRNAs (i.e. presence vs absence) to predict primary sites using a 3-fold cross-validation framework. Oral cavity, oropharynx, and larynx primary subsite classification achieved area-under-the-curve (AUC) consistently over 0.75 (**Figures 1D, S1B**). Furthermore, oncRNA profiles can effectively stratify tumors by anatomical site, namely oral cavity, oropharynx, hypopharynx, and larynx. (**Figures 1E, S1C-F**). Our results indicate that similar to gene expression profiles, digital oncRNA fingerprints can be utilized to accurately classify and predict primary sites for HNSCC.

### Validation of Oral Cavity oncRNA Signatures in an Independent Institutional Cohort

Given that oral cavity squamous cell carcinoma (OCSCC) is the most prevalent subsite of HNSCC worldwide and possesses well-established murine models for functional validation, we prioritized the investigation of oncRNA profiles in patients with OCSCC. To validate the signatures identified in public datasets, we performed oncRNA profiling on an independent institutional cohort composed primarily of patients with advanced-stage OCSCC. We confirmed that the specific repertoire of oral cavity oncRNAs identified *in silico* is robustly detectable in solid tumors from this clinical cohort. Importantly, these transcripts displayed high specificity; they were not enriched in patients with different primary cancers of the head and neck region, reinforcing their utility as precise molecular fingerprints for OCSCC (**Figures 2A, S2A**).

**Figure 2.**
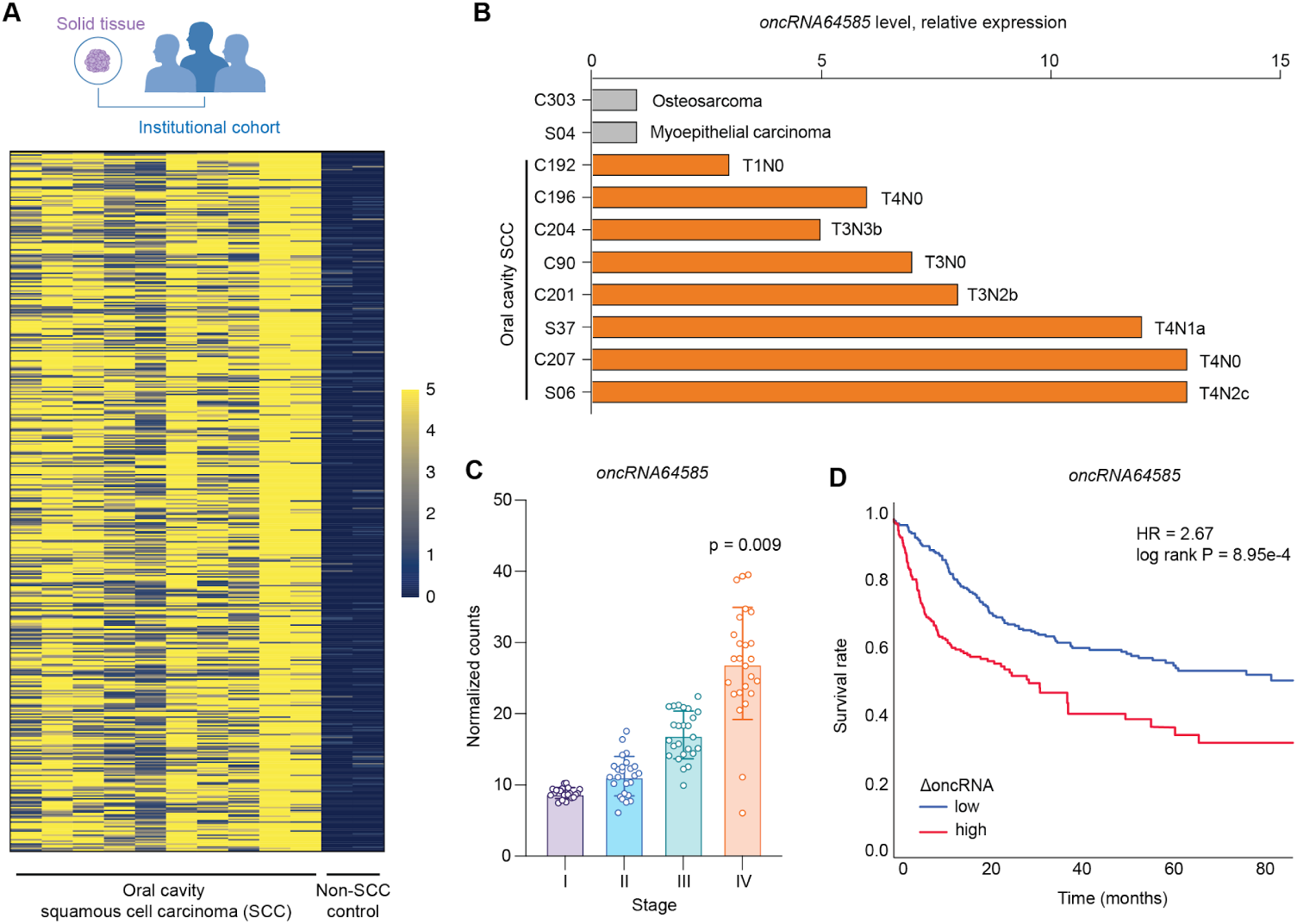
oncRNA64585 expression level correlates directly with disease progression and prognosis. **(A)** Heatmap showing expression levels of top oncRNAs enriched in solid tumors from an institutional retrospective cohort (n = 10). C303 and S04 are non-squamous cell carcinoma cancer patients serving as negative controls. **(B)** Bar-graph showing expression levels of *oncRNA64585* in a cancer type- and stage-dependent manner. **(C)** Boxplots showing higher levels of oncRNA64585 are associated with more advanced disease in the TCGA database. P-value performed with Wilcoxon rank test. **(D)** Survival plot comparing patients with higher *oncRNA64585* expression (2-fold higher than the mean) versus lower expression (2-fold lower than the mean).

### *oncRNA64585* Expression Correlates with Advanced Disease and Poor Prognosis

Having validated the OCSCC oncRNA signature, we sought to identify transcripts with potential functional roles in disease progression. Among the validated oral cavity oncRNAs, *oncRNA64585* emerged as a priority candidate due to its consistent enrichment across samples (**Figure S1C**). Analysis of clinical demographics from the TCGA cohort revealed that elevated oncRNA64585 expression is significantly associated with higher tumor grade (**Figure 2B**) and more advanced disease stage (**Figure 2C**). Furthermore, high expression of oncRNA64585 correlated with worse overall survival (**Figure 2D**). Of note, most of the patients in the TCGA cohort presented with primary oral cavity (tongue) and advanced stage (**Figures S2B and C**). Additionally, specificity of this association was confirmed by analyzing separate OCSCC-enriched oncRNAs, which showed no correlation with patient survival (**Figure S2D**). These strong clinical associations suggest that *oncRNA64585* is not merely a passenger transcript but potentially serves a functional role in tumorigenesis, warranting further mechanistic investigation.

### Cross-Species Analysis Identifies *oncRNA64585* as a Conserved Regulator of Tumor Progression

To investigate the functional relevance of our candidates, we utilized the matched pair of syngeneic Murine Oral Cavity (MOC) carcinoma models: MOC1 (indolent phenotype) and MOC2 (aggressive phenotype) ^18,19^. OncRNA profiling in these models revealed a subset of oncRNAs that are shared between human and mouse (**Figure 3A**). Notably, we identified *oncRNA64585* as one of the most highly abundant oncRNAs conserved across both species (**Figure 3B**). Importantly, we detected *oncRNA64585* in both MOC1 and MOC2 lines, but not in the breast cancer cell line 4T1, further highlighting its specificity to OCSCC (**Figure S3A**).

**Figure 3.**
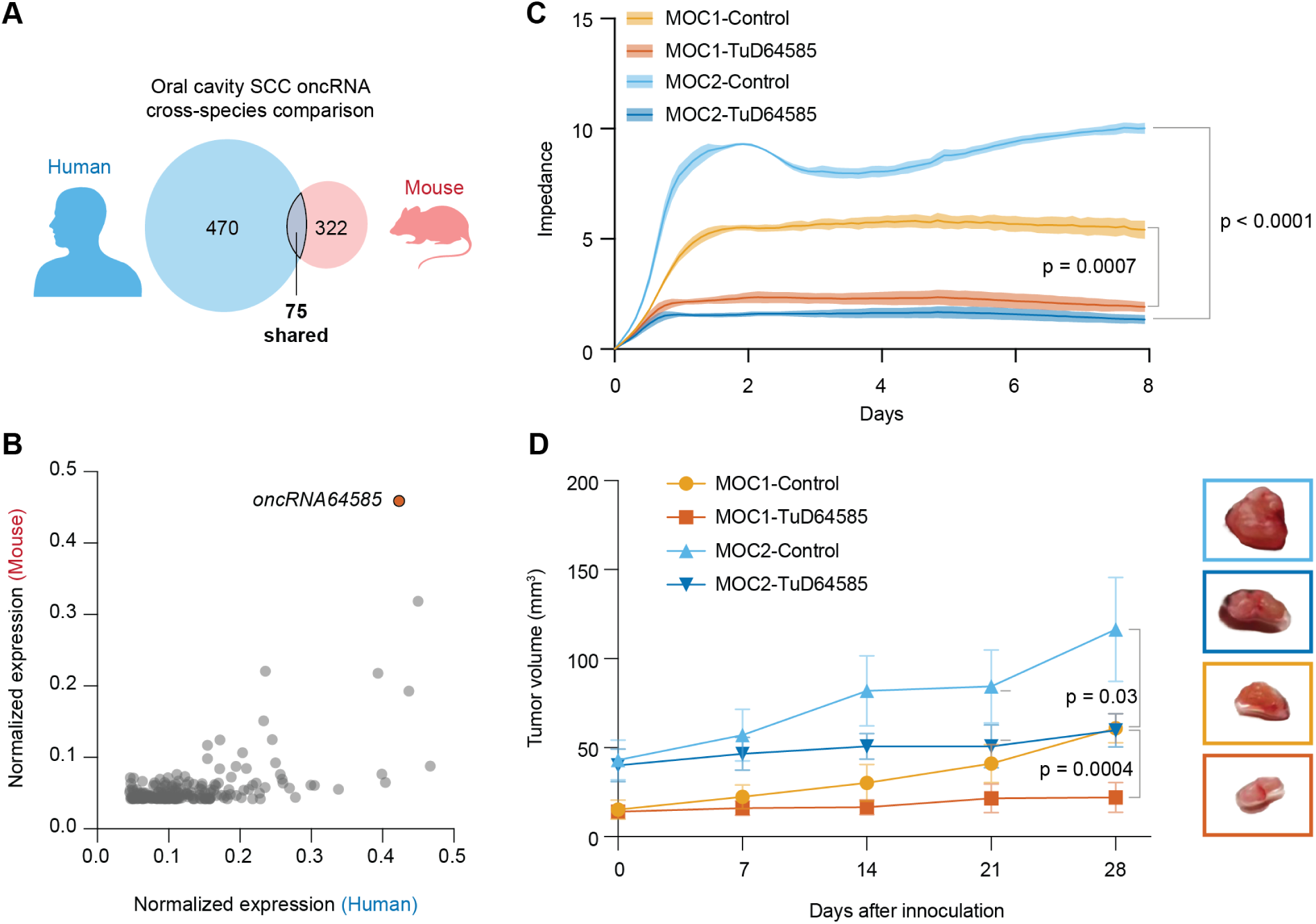
*oncRNA64585* expression promotes tumor growth. **(A)** Venn diagram showing overlap between human and mouse OCSCC oncRNAs. **(B)** Scatter plot showing correlation in terms of expression levels (in count-per-million) of oncRNA64585 between human and mouse. **(C)** *In vitro* growth curves of MOC1 and MOC2 cells, with either control or anti-oncRNA64585 Tough Decoy (TuD) vectors, measured as a function of impedance by Xcelligence Assay over two weeks (*n* = 6) . **(D)** *In vivo* tumor growth of MOC1 and MOC2 cells, with either control or anti-oncRNA64585 TuD (*n* = 5).

Loss-of-function experiments utilizing anti-sense Tough Decoy (TuD) constructs targeting *oncRNA64585* (**Figures S3B-C**) resulted in a significant reduction in *in vitro* proliferation for both cell lines (**Figure 3C**). Notably, the growth of the aggressive MOC2 line was blunted to levels comparable to the indolent MOC1 line, underscoring the importance of this oncRNA in tumor progression. Furthermore, mice inoculated with TuD-*oncRNA64585*-expressing MOC1 and MOC2 cells exhibited >50% reduction in tumor volume at day 28 compared to controls, recapitulating our *in vitro* findings (**Fig. 3D**). Taken together, these findings strongly implicate *oncRNA64585* as a direct regulator of tumor growth and disease progression.

### *oncRNA64585* Controls Tumor Growth via Chromatin-Mediated Regulation of Cell Cycle and Apoptosis Genes

To investigate the mechanism by which *oncRNA64585* drives tumor growth, we performed transcriptional profiling on ex-vivo tumors harvested at endpoint (day 28). TuD-mediated knockdown of *oncRNA64585* induced marked global transcriptional remodeling, with over 3,000 differentially expressed genes identified (**Figures 4A** and **S4A**). Pathway analysis linked these genes to key oncogenic hallmarks, including interferon signaling and epithelial-mesenchymal transition (**Figure S4B**). To identify core drivers of tumorigenesis, we isolated genes with conserved expression changes across both MOC1 and MOC2 models following *oncRNA64585* depletion. We identified 282 up-regulated and 118 down-regulated genes; notably, these perturbations were more pronounced in the aggressive MOC2 line (**Figure 4B**), consistent with the more dramatic tumor suppression observed in vivo. Functional enrichment analysis revealed that downregulated genes were primarily involved in cell cycle regulation (**Figure 4C**, blue), while upregulated genes were associated with pro-inflammatory and pro-apoptotic pathways (**Figure 4C**, red). ChIP-X enrichment analysis implicated the Polycomb Repressive Complex 2 (PRC2) components SUZ12, RNF2, and MTF2 as upstream regulators (**Figure S4C**). These results suggested that *oncRNA64585* promotes oncogenesis by modulating chromatin state.

**Figure 4.**
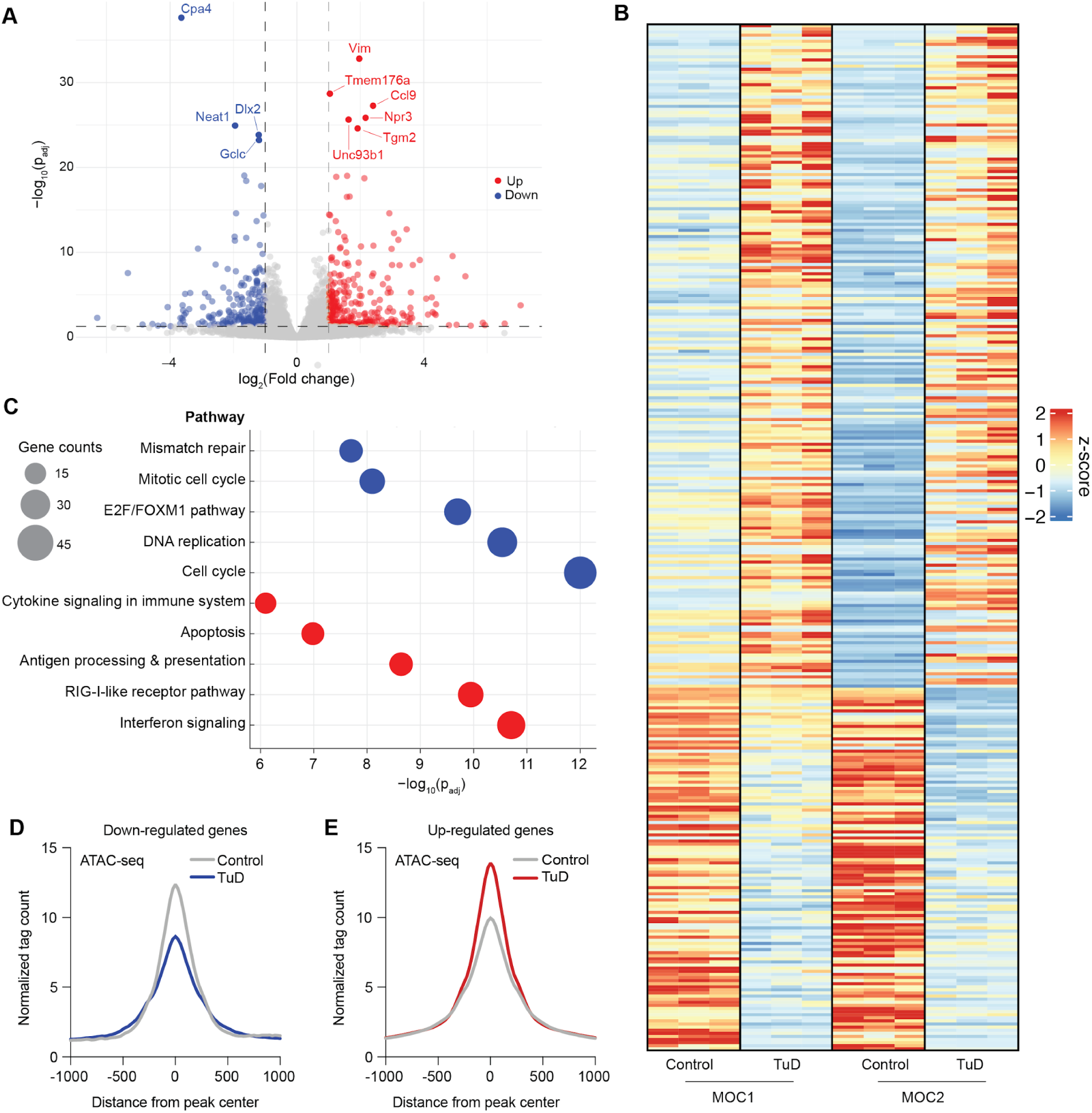
Global transcriptional and chromatin accessibility disruption as a result oncRNA64585 loss-of-function. **(A)** Volcano plot showing over 3000 differentially expressed genes between oncRNA64585 knock-down compared to control in MOC1 cell lines (FDR < 0.05, |log2FC| >=1). **(B)** Heatmap showing shared differentially expressed genes between MOC1 and MOC2 with control or oncRNA64585 KD. **(C)** Gene ontology analysis of 282 up-regulated (red) and 118 down-regulated (blue) as identified from (B). **(D)** and **(E)** Average density profiles of ATAC seq data showing mean coverage (n = 2 biological replicates) at all identified peaks associated with genes down-regulated or up-regulated as a function of oncRNA64585 KD, respectively.

To test this hypothesis, we performed ATAC-seq on control and *oncRNA64585-*knockdown MOC2 cells. Crucially, we observed decreased chromatin accessibility (more compact state) within 100kb of the promoters of downregulated genes (**Figures 4D** and **S4D**). Conversely, upregulated genes were associated with increased chromatin accessibility (**Figures 4E** and **S4E**). We next sought to define the molecular mechanism by which oncRNA64585 regulates gene expression. RNA immunoprecipitation (RIP) assays revealed significant enrichment of oncRNA64585 in SUZ12 immunoprecipitates compared with IgG controls, indicating a direct or stable association with the PRC2 complex (**Figure S4F**). These results support a model via which *oncRNA64585* acts as a regulatory RNA that modulates PRC2 activity to reprogram chromatin accessibility and gene expression.

### Pre-Diagnostic Detection of Oral Cavity Cancer via AI-Guided Analysis of Plasma oncRNAs

Early detection and risk stratification are paramount in HNSCC. In HPV-negative disease especially, the lack of reliable biomarkers negatively impacts overall survival. Stratifying high-risk individuals before clinical manifestation could enable timely intervention, limit disease progression, and reduce treatment-associated morbidity. We previously established that while the majority of oncRNAs are retained intracellularly, a distinct fraction is actively secreted into the circulation, rendering them robust candidates for liquid biopsy applications in minimal residual disease detection^12,13^ as well as disease detection^15,27^. To see if HNSCC-associated oncRNAs could be similarly utilized as biomarkers from liquid biopsies, we first validated detection feasibility by profiling tumor-matched plasma samples from our institutional cohort (previously shown in Figure 2A). We confirmed that a significant portion of solid-tumor oncRNAs is detectable in cell-free plasma (**Figure S5A**), exhibiting expression profiles highly concordant with their matched solid tumors (**Figure 5A**).

**Figure 5.**
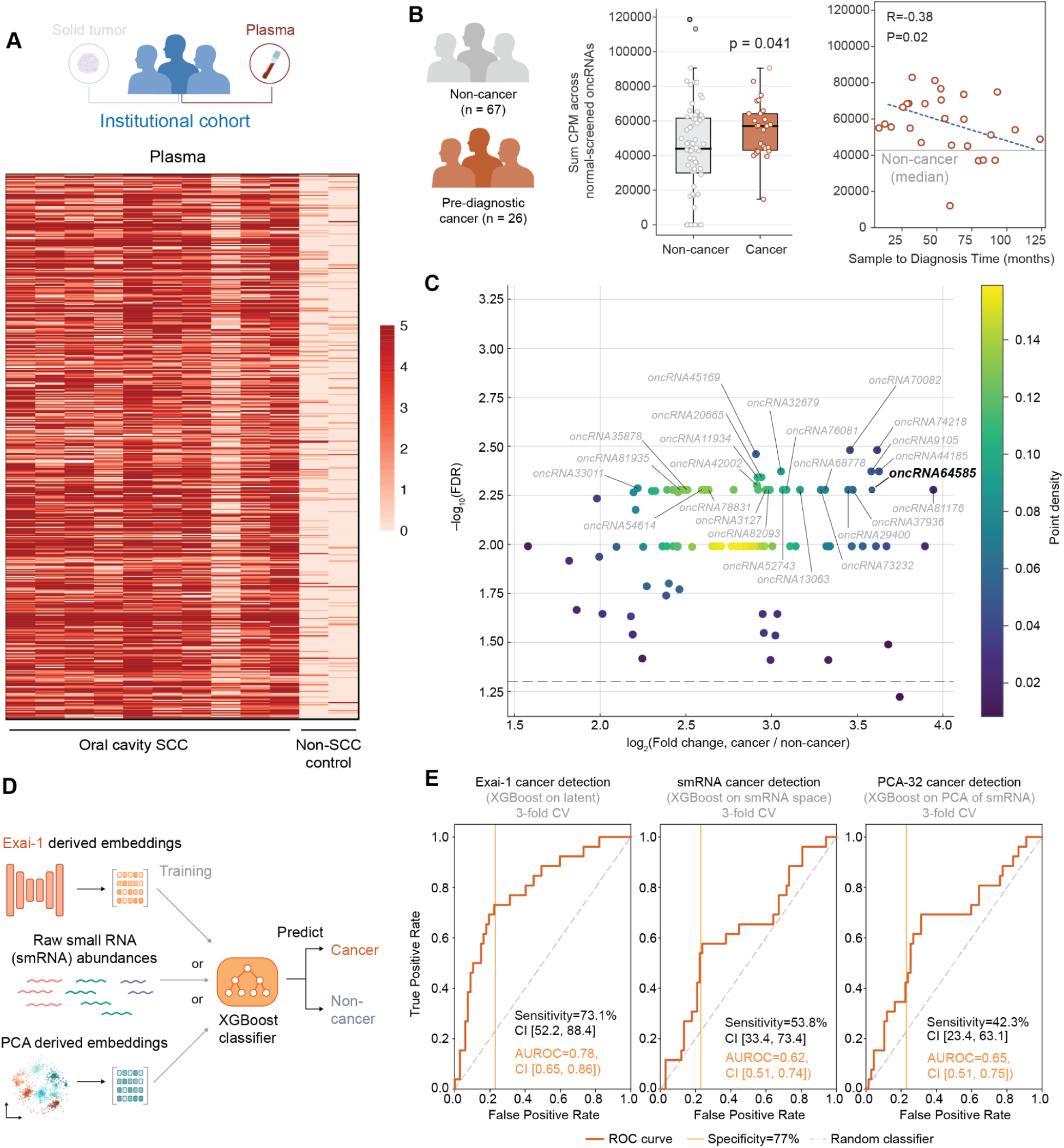
OncRNA profiles can predict cancer prior to diagnosis using liquid biopsies. **(A)** Heatmap showing corresponding oncRNA profile of matched tumor-plasma samples. **(B)** Boxplots showing total sum expression level of all oncRNAs between 26 pre-diagnostic cancer and 67 non-cancer samples. P-value was calculated with Mann-Whitney test. The scatter plot depicts this aggregated oncRNA burden as a function of time from sample to diagnosis. Shown are the Pearson correlation and the associated *p*-value (one-sided). **(C)** Dot plot showing the differential expression of the top 100 oncRNAs previously identified from a combination of TCGA, CPTAC-3, and institutional cohort in this independent pre-diagnostic cohort of 93 patients. **(D)** Schematic of Exai-1 predictive model. **(E)** Area Under the ROC Curve (AUC) plots comparing performance from different machine learning algorithms on stratifying pre-diagnostic cancer from non-cancer patients.

To evaluate the predictive value of circulating oncRNAs, we analyzed pre-diagnostic biobanked plasma samples from a prospectively collected cohort. We identified 26 patients who subsequently developed oral cavity cancer (tongue primary) and 67 age-, gender-, and smoking-matched controls who remained cancer-free (**Table 1**). Notably, the median lead time to clinical diagnosis was over 4 years (50.2 months). As expected, global oncRNA burden, defined as the total sum of abundances across all annotated HNSCC oncRNAs, was significantly elevated in pre-diagnostic cancer samples compared to controls (**Figure 5B**). Similarly, this aggregate oncRNA burden score was significantly anti-correlated with the sample to diagnosis time among the cancer cohort (**Figure 5B**). Thus, circulating oncRNA burden captures a cancer-specific signal, detectable well in advance of clinical diagnosis.

**Table 1.** Characteristics of Cases and Matched Controls.

| **Characteristic** | **Case**<br>**N = 26<sup>1</sup>** | **Control**<br>**N = 67<sup>1</sup>** |
| --- | --- | --- |
| **Sex** |  |  |
| Male | 14 (54%) | 32 (47%) |
| Female | 12 (46%) | 36 (53%) |
| **Smoking History** |  |  |
| Never | 14 (54%) | 32 (47%) |
| <10 pack-years | 5 (19%) | 15 (22%) |
| ≥10 pack-years | 7 (27%) | 21 (31%) |
| **Age at Blood Draw**<br>**(years)** | 66.5 [56.0, 73.0] | 62.3 [52.3, 66.0] |
| **Time from sampling to**<br>**diagnosis (months)** | 50.2 [25.4, 81.7] | — |
| <sup>1</sup> n (%); Median [Q1, Q3] |  |  |

Next, we focused on the top 100 oncRNAs that were previously identified from a combination of public datasets (TCGA, CPTAC-3) as well as our solid tumor-plasma matched cohort. These oncRNAs, including *oncRNA64585*, were consistently upregulated in pre-diagnostic plasma samples (**Figure 5C**), and successfully differentiated future cancer patients from controls (**Figure S5B**).

Finally, we sought to maximize detection sensitivity by leveraging Exai-1, a multi-scale cell-free RNA foundation model we recently introduced ^16^. Pre-trained on over 13,000 liquid biopsy profiles, Exai-1 learns to map cell-free small RNA sequencing data, which is zero-inflated by nature due to sparse and stochastic sampling of low-abundance circulating RNAs, into a robust, 32-dimensional “latent representations” that capture biological signal while suppressing technical variation. Here, we applied a transfer learning framework to leverage Exai-1’s learned representations for few-shot classification on our HNSCC cohort. We compared classifiers trained on these Exai-1 embeddings against those trained on standard feature spaces: raw small RNA (smRNA) abundances and PCA-derived embeddings. Classifiers utilizing Exai-1 latents consistently achieved higher discrimination between pre-diagnostic HNSC plasma samples and controls (**Figure 5D**). While PCA provided a reasonable baseline, raw smRNA features underperformed, likely reflecting their sensitivity to assay noise. By contrast, the Exai-1 latent space, which incorporates structural and sequence context for thousands of oncRNAs, resulted in a superior area under the ROC curve (AUROC) (**Figure 5E**). In a 3-fold cross-validation scheme, Exai-1-based classifiers achieved a 73% sensitivity at 77% specificity. These results are particularly notable given the challenging pre-diagnostic setting, where samples were collected years prior to clinical diagnosis (median >4 years). Given these extensive lead times, tumor burden is likely minimal if not entirely absent in some samples, presenting a significant hurdle for blood-based detection. These findings demonstrate that transferring learned representations from our large-scale foundation model enables robust cancer detection even when the biological signal from oncRNA is highly attenuated.

## DISCUSSION

HPV-negative HNSCC remains one of the most challenging malignancies to manage due to its aggressive biology and lack of early detection strategies. Although liquid biopsies have gained momentum, their utility in this cohort has been constrained. Circulating tumor DNA (ctDNA) demonstrates limited sensitivity in early-stage disease and often necessitates tumor-informed designs that are feasible in surveillance but challenging for screening ^20–22^. Similarly, methylation signatures and mutation panels face technical barriers regarding limit-of-detection and sequencing depth ^23–26^. Our study introduces orphan non-coding RNAs (oncRNAs) as an orthogonal, high-specificity biomarker class that addresses several of these limitations.

By leveraging small-RNA transcriptomic data from TCGA and CPTAC-3, we described the comprehensive landscape of oncRNAs in HPV-negative HNSCC, demonstrating their ability to stratify tumors by site, predict cancer from plasma, and functionally promote oncogenesis. These findings build upon our foundational work characterizing oncRNAs as cancer-specific, unannotated transcripts ^12,13,27^. Because treatment strategies for head and neck cancers vary significantly by subsite, the ability to utilize oncRNA fingerprints to predict the tumor’s anatomical origin enhances diagnostic accuracy—particularly in cases of unknown primary. Clinically, our profiling of pre-diagnostic plasma samples demonstrated that oncRNA signatures could predict future oral cavity cancer with high accuracy. The ability to detect these signals years prior to diagnosis (median lead time >4 years) highlights the transformative potential of oncRNAs for early risk assessment in asymptomatic populations.

Moving from association to function, we identified oncRNA64585 as a key driver of tumor growth. Using syngeneic murine models, we demonstrated that knockdown of oncRNA64585 significantly impaired in vitro proliferation and in vivo tumor progression. Mechanistically, RNA-seq and ATAC-seq analyses revealed that this transcript regulates cell cycle and apoptosis programs via the modulation of chromatin accessibility through the PRC2 complex. These data suggest that oncRNA64585 acts as a cancer-emergent neo-regulator of the epigenome, exploiting cryptic promoter activation to drive tumorigenesis.

Our results also highlight the advantages of applying foundation models such as Exai-1 to cfRNA datasets in oncology. Exai-1 enables the construction of accurate diagnostic classifiers even in settings where sample size is modest, such as the present HNSCC plasma cohort. Other feature representations we evaluated did not approach the performance achieved with Exai-1, underscoring the value of model pre-training on large and diverse cfRNA corpora. Exai-1 is a pre-trained foundation model and is not tailored to a particular cancer type. It is instead designed to capture the breadth of cfRNA biotypes relevant to liquid biopsy, relying heavily on oncRNAs to capture cancer-relevant signals. Together, these findings support the utility of foundation models such as Exai-1 as practical tools for building diagnostic classifiers across cancers where data availability is limited.

Collectively, our results propose a new paradigm in precision oncology: one that incorporates cancer-emergent RNA species to refine detection and stratification. OncRNAs represent an underexplored molecular axis in HNSCC, and their integration into clinical strategies holds the potential to shift the management of HPV-negative disease. Furthermore, our findings intersect with a rapidly evolving therapeutic landscape. Recent approvals of RNA-targeted therapies—including antisense oligonucleotides (ASOs) and small RNA inhibitors—demonstrate the clinical feasibility of this modality. Our in vivo success with Tough Decoy inhibitors serves as a proof-of-concept that targeting specific oncRNAs, such as *oncRNA64585*, is a viable therapeutic strategy.

## METHODS

### Public cohort analysis (TCGA-HNSC and CPTAC-3)

#### Data acquisition and sample selection

Bulk miRNA-seq data and clinical annotations for TCGA-HNSC and CPTAC3 were obtained from the Genomic Data Commons (GDC). We retained primary tumor samples with complete site and other demographics. All analyses reported here used human genome build hg38 for consistency across cohorts.

#### Alignment and expression quantification

To quantify oncRNA loci, we used a custom GTF consisting of short, unannotated, tumor-enriched loci curated in prior discovery analyses. Counts for these loci were generated in parallel to gene-level counting using the same coordinate system to ensure cross-cohort comparability.

#### Normalization, filtering, and batch awareness

Raw counts were normalized using standard methods suitable for between-sample comparisons (library-size normalization with variance-stabilizing/regularized-log transforms for visualization; model-based normalization for differential testing). Features with extremely low abundance across the cohort were filtered a priori to reduce multiple-testing burden. When combining TCGA with CPTAC, we accounted for dataset-level effects by including cohort as a covariate in linear models; visualization panels (e.g., PCA) were generated on normalized values with batch-aware scaling for plotting. Where appropriate, differential analyses controlled for relevant clinicopathologic covariates (e.g., anatomical site). Multiple-testing corrections used Benjamini–Hochberg false-discovery rate.

#### Stratification and replication

To assess site-specific structure, we performed PCA/hierarchical clustering on normalized oncRNA profiles with Euclidean distance and Ward’s linkage. Cross-cohort replication of OCSCC-enriched oncRNAs was evaluated by concordance of effect direction and significance in TCGA vs. CPTAC, using identical filtering and modeling criteria in each cohort to avoid thresholding artifacts.

#### Cell Culture

MOC1 and MOC2 murine oral cavity carcinoma cell lines were obtained cultured from Sigma Aldrich (SCC469, SCC470). Cultures were grown in DMEM supplemented with 10% FBS and penicillin-streptomycin according to manufacturer’s recommendations.

#### In vitro cell proliferation assay

For impedance-based growth measurements, xCELLigence E-Plate 96 microelectrode plates (Agilent/ACEA Biosciences) were pre-equilibrated with complete medium (100 µL per well) for 30 min at room temperature to obtain background readings. Cells were resuspended in complete medium, counted with a hemocytometer/automated counter, and seeded at 10,000 cells per well in a final volume of 200 µL seeding density optimized to yield a Cell Index (CI) of ∼0.3–0.5 within 16–24 h. Edge wells were filled with sterile PBS or medium to minimize evaporation. Plates were placed in the xCELLigence RTCA MP/DP instrument housed inside a standard CO₂ incubator, and CI was recorded every 15 min for the first 6 h and every 30–60 min thereafter for the duration of the assay.

#### Stem-loop RT–qPCR

Assay was designed and implemented using the Thermo Fisher TaqMan™ MicroRNA stem-loop architecture (custom design), with the manufacturer’s positive control used as a reference for assay setup and performance benchmarking. Amplification used a forward primer to the target and a universal reverse primer complementary to the stem-loop backbone; detection employed a short hydrolysis probe to ensure adequate Tm for the small amplicon. Reverse transcription (stem-loop RT) on total RNA (tissue or plasma eluate), followed by qPCR (95 °C 10 min; 40 cycles of 95 °C 15 s, 60 °C 60 s). NTC and no-RT controls were included in every run. All primer oligos are listed in **Table 2**

**Table 2.**
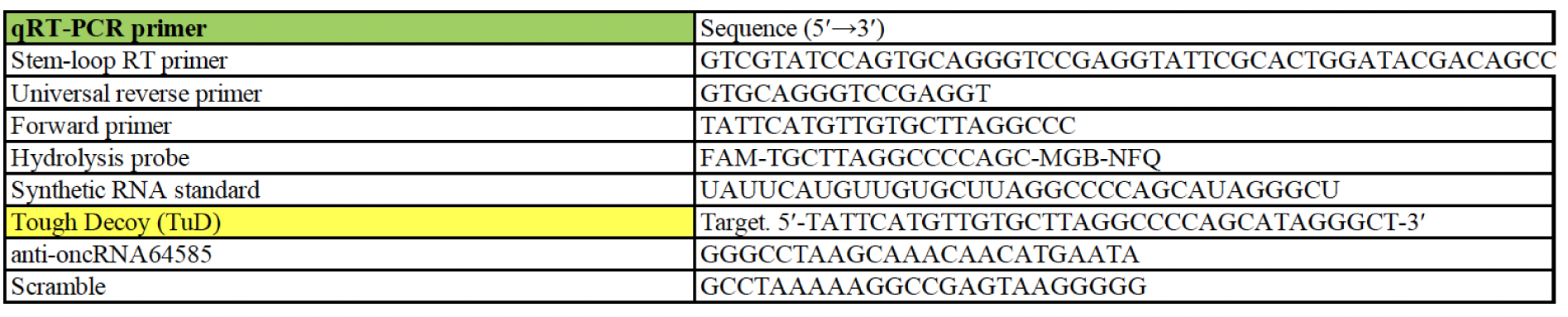
Oligonucleotide sequences used in this study.

#### UV Crosslinking Immunoprecipitation of Suz12-bound oncRNA

MOC1 cells were UV irradiated (400 mJ cm−2) before lysis with buffer (1×PBS, 0.1% SDS, 0.5% sodium deoxycholate, and 0.5% Igepal CA-630) supplemented with 0.2U/μl SUPERase·In RNase Inhibitor (Thermo Fisher) and 1x Halt protease inhibitor cocktail (Thermo Fisher). Lysates were then treated with 0.1 U/μl RQ1 RNase-free DNase (Promega) at 37°C for 10 min and cleared by centrifugation. Then, Suz12-RNPs were immunoprecipitated with a Suz12-specific antibody (Cell Signaling #3737) bound by protein A dynabeads (Life Technologies) at 4°C for 2 hr. In parallel, IgG isotype control (Proteintech #30000-0-AP) was included. The immunoprecipitants were washed twice with lysis buffer, twice with high salt buffer (5×PBS, 0.1% SDS, 0.5% sodium deoxycholate, and 0.5% Igepal CA-630) and twice with PNK buffer (50 mM Tris-HCl, pH 7.4, 10 mM MgCl2, and 0.5% Igepal CA-630). 4 mg/ml of Proteinase K in PK buffer (50 mM Tris-HCl, pH 7.5, 50 mM NaCl, and 10 mM EDTA) was added to the beads or input lysates and shaken at 55°C for 20 min. Immunoprecipiated RNAs were then extracted using Trizol reagent (Thermo Fisher) followed by phase-seperation with chloroform by centrifugation at 4°C for 10 min. The upper aqueous phase was then transferred to a new tube and RNAs were precipitated with 100% ethanol, 0.3 M NaAC and glycoblue co-precipitant (Thermo Fisher) for 1 hr at -80°C

#### Tough Decoy (TuD) inhibitor targeting oncRNA

A TuD bearing two antisense binding sites (BS) to the oncRNA was engineered, each with a 2-nt bulge at positions 10–11 (relative to the target) to prevent Ago2 cleavage while preserving high-affinity binding. Binding windows were chosen within the 34-nt target to avoid extreme GC or homopolymer runs. All oligo sequences for TuD are listed in **Table 2**.

Oligos were synthesized (5′-phosphorylated), annealed, and inserted into a standard TuD hairpin scaffold driven by a U6 promoter within a lentiviral transfer plasmid with a pLKO backbone using Gibson assembly per vector specifications. The complete TuD cassette (BS1 and BS2 on opposing arms) and the scramble control were sequence-verified.

#### Lentiviral production

HEK293T cells (70–80% confluence) were co-transfected with the transfer plasmid (U6-TuD) and **t**hird-generation packaging plasmids (pMDLg/pRRE) plus VSV-G envelope (pMD2.G) using Lipofectamine 2000. Medium was replaced after 6–8 h. Viral supernatants were harvested at 48 h and 72 h, clarified (0.45 µm), and optionally concentrated (PEG or ultracentrifugation). Viral titer was estimated by p24 ELISA or functional transduction of indicator cells.

#### Transduction and selection

Target cells were transduced at MOI 0.3–1 (single-copy bias) in the presence of polybrene 4–8 µg/mL; spinoculation was used as needed. At 24 h, medium was refreshed; selection began at 48–72 h using the vector’s antibiotic puromycin 2 µg/mL.

#### In Vivo Tumor Growth Assay

TuD-expressing MOC1 and MOC2 cells (5 × 10⁶ cells per flank) were injected subcutaneously into C57BL/6 mice (n = 5 per condition). Tumor volumes were measured every 7 days with digital calipers and calculated using the formula V = (length × width²)/2. Mice were sacrificed at day 28, and tumors were harvested for downstream sequencing. Experiments were conducted under IACUC-approved protocols.

#### Sequencing library preparation and analysis

For oncRNA, Total RNA (DNase-treated as needed) was prepared using the SEQuoia Complete RNA Library Preparation Kit following the manufacturer’s instructions. Briefly, the workflow incorporates rRNA reduction, controlled enzymatic fragmentation, random-primed first-strand synthesis, and dUTP-based second-strand synthesis to yield stranded libraries. After end-repair/A-tailing, unique dual-indexed adapters were ligated; libraries were PCR-amplified for the minimal recommended cycles, bead-purified, QC’d by fragment analysis and fluorometric quantification, pooled, and sequenced on an Illumina platform (paired-end, 2×100bp). Demultiplexed reads were trimmed and aligned to hg38 (or mm10 for murine samples) with reverse-strand counting. Where UMIs were present, PCR duplicates were collapsed in a UMI-aware step prior to quantification. Gene-level counts used a GENCODE GTF; oncRNA loci were quantified with a custom GTF. Downstream normalization and modeling followed the public-cohort pipeline above.

For transcriptomic profiling, tissues were rinsed once with ice-cold PBS and lysed directly with Qiagen RNAeasy kit (Cat 74104) . Total RNA was extracted following the manufacturer’s protocol, including on-column DNase I treatment where applicable. RNA quantity and purity were assessed by NanoDrop (A₂₆₀/A₂₈₀ and A₂₆₀/A₂₃₀), and integrity was measured by Agilent 2100 Bioanalyzer; only samples with RIN ≥ [8.0] proceeded to library preparation.

RNA processing, library construction, and sequencing were performed by Novogene (Sacramento, USA) following their standard mRNA-seq pipeline. Libraries were size-selected to an average insert of ∼200–300 bp, quantified by qPCR, pooled equimolarly, and sequenced on an IlluminaNovaSeq 6000 platform to generate 150-bp paired-end reads. Each sample achieved a minimum depth of ≥20 million paired reads

Read processing, alignment, and quantification: Adaptor trimming and low-quality base removal were performed by Novogene using cutadapt with default stringency; in-house QC used FastQC to verify per-base quality and duplication metrics. Reads were aligned to the human reference genome mm10 using STAR (v2.7.2) ^28^ with parameters optimized for splice-aware mapping (e.g., --twopassMode Basic, --outFilterMismatchNmax 999, --outFilterMismatchNoverReadLmax 0.04, --alignSJDBoverhangMin 1). Gene-level raw counts were obtained via STAR’s --quantMode GeneCounts (unstranded/stranded flag set to match library chemistry) using the corresponding GTF.

#### Differential gene expression analysis

Downstream analysis was conducted in R (4.3.1) with DESeq2 (v1.40.2)^29^. Low-count genes were filtered a priori (keep genes with ≥10 counts in ≥2 samples). Size factors were estimated by the median-of-ratios method, and dispersion parameters were fit using DESeq2 defaults. Unless specified otherwise, contrasts tested the effect of [microplastics dose or condition] vs vehicle control using Wald tests with Benjamini–Hochberg false discovery rate (FDR) correction. Genes were considered differentially expressed at FDR < 0.05 with an absolute log₂ fold change (|log₂FC|) threshold of ≥ 1 when indicated. For effect-size shrinkage in visualization and ranking, lfcShrink with the apeglm method was applied. Variance-stabilizing transformation (VST) or regularized log (rlog) counts were used for principal component analysis (PCA) and clustering.

#### Pathway and gene-set enrichment

Pathway analysis was performed with Enrichr (using differentially expressed gene (DEG) sets (up- and down-regulated tested separately unless noted). The following libraries were queried: GO Biological Process 2025, ChEA2022. En enrichment significance was evaluated using Enrichr’s combined score and adjusted p-values (Benjamini–Hochberg). Where indicated, gene-set enrichment analyses were corroborated by over-representation tests in clusterProfiler using the same background universe (all tested genes).

#### ATAC-seq library generation and data processing

ATAC-seq libraries were prepared by Novogene. Briefly nuclei were isolated and tagmented using Nextera reagents, followed by PCR amplification and library preparation. Sequencing data were aligned to mm10 genome using Bowtie2 (v2.2.6) ^30^ with the following parameters -p 20 -N 1. Mapped reads were processed into tag counts, filtered for PCR duplicates, corrected for read-depth bias, and fragment lengths for each biological replicate with HOMER ^31^ functions makeTagDirectory -tbp 1 -fragLength 150 -totalReads 2e7. For peak calling, findPeaks function was used with parameter -size 200 to resize all peaks to a uniform size of 200bp. Genome browser tracks were generated with HOMER function makeUCSCfile -bigWig -fsize 1e20 and corrected for variations in read-depth with -norm e7. Average peak profiles were made with function annotatePeak with resulting coverage-by-nucleotide profile visualized with Prism Graphpad.

#### Data visualization

Heatmaps were generated with “pheatmap” (v1.0.12) from z-scored VST-normalized expression values. Rows (genes) and columns (samples) were clustered using 1 – Pearson correlation as the distance metric and complete linkage (row/column scaling as indicated in figure legends). Row annotation tracks show gene-set membership or pathway labels where applicable. Additional plots (PCA, volcano plots, MA plots) were created with “ggplot2 “(v3.5.2) and EnhancedVolcano (v1.18.0) using consistent aesthetic parameters across figures. All analysis code (R scripts and STAR command lines) is available upon request on publication. Raw and processed RNA-seq data will be deposited in GEO database.

### Pre-diagnositic plasma oncRNA profiling and cancer prediction

#### Sample acquisition

Plasma samples of pre-diagnostic head and neck cancer patients were obtained from the Mass General Brigham Biobank. Our initial goal was to have a 1:3 (cancer:non-cancer) ratio to help with a) obtaining adequate number of non-cancer patients to establish baseline level of plasma oncRNAs and 2) adequate randomization for machine learning training and classification. We were able to identify 26 primary oral cavity squamous cell carcinoma (oral tongue) and 68 age-, gender-, smoking-matched controls. One of out of 68 control samples did not meet quality control criteria prior to sequencing.

#### Cohorts and data processing

Raw counts were normalized to counts-per-million (CPM) per sample, and log(1+CPM) was used where indicated to stabilize variance.

#### Summary score

To demonstrate signal without machine learning, we first removed oncRNAs with evidence of background expression in normals. Ten non-cancer samples were randomly selected (seed = 42). Any oncRNA with CPM ≥ 1 in any of these 10 normal was excluded. All subsequent “no-ML” summaries were computed on the retained set. (A K-of-10 and threshold-tunable variant was used in sensitivity analyses.) Per sample, the sum of CPM across retained oncRNAs was calculated.

#### Confirmatory Machine learning model

Additionally, we performed weighted ranked sum using differentially expressed oncRNA previously identified. For each retained oncRNA we computed mean CPM in cancer/non-cancer and prevalence (fraction CPM ≥ 1). Weights were w_i = max(0, log2[(μ_i^c+1)/(μ_i^n+1)]) × max(0, p_i^c − p_i^n). Features were ranked by weight; the per-sample score was Σ_{TopN} w_i · log(1+CPM_i). We report N = 100 (with N swept 50–500 in sensitivity analyses).

We fitted linear classifiers (logistic regression, linear SVM; class-weighted) on log(1+CPM) features with standardization and simple feature budgets (Top-K by univariate enrichment/coefficients). Internal stratified evaluation produced ROC/PR curves, and decision thresholds were chosen to satisfy ≥90% precision.

#### Reproducibility

Analyses were conducted in Python (pandas, numpy, scikit-learn, matplotlib) with fixed random seed (42). All parameters (CPM thresholds, K-of-10 rule, and N) are reported above and were varied only in prespecified sensitivity analyses.

#### Cancer detection benchmarking with Exai-1 latent space, raw smRNA features, and PCA embeddings

We leveraged Exai-1, a multi-modal generative transformer foundation model trained and evaluated on over 13,000 cell-free RNA (cfRNA) profiles from plasma and serum samples. Exai-1 integrates cfRNA sequence embeddings with small RNA (smRNA) abundance profiles to learn biologically meaningful latent representations that capture co-expression structure, denoise technical variation, and improve cross-biofluid generalization. Exai-1 models sequence and structure 7,349 cfRNAs, including 4,558 oncRNAs, 761 yRNAs, 716 miRNAs, 704 tRNAs, and 610 snoRNAs. The model uses a variational transformer architecture augmented with contextual special tokens (e.g., cancer status, biofluid type) and outputs a 32-dimensional latent embedding per sample. To generate PCA features, smRNA profiles were z-scored across samples and reduced to 32 principal components (scikit-learn v1.4, full SVD solver). Full model details and training procedures are described in the original Exai-1 publication (Karimzadeh et al., 2025, bioRxiv).

#### Study cohort and feature sets

For this study, we analyzed a set of plasma samples from patients with head and neck squamous cell carcinoma (HNSC) and non-cancer controls. For each sample, three different feature representations were evaluated for cancer detection:

- smRNA abundance space — normalized log1p counts-per-million (CPM) values across 7,349 cfRNAs used in Exai-1 training.
- PCA space — the top 32 principal components derived from the z-scored log1p CPM smRNA abundance matrix, chosen to match the dimensionality of Exai-1 latents.
- Exai-1 latent space — 32-dimensional embeddings obtained directly from the trained Exai-1 encoder.

#### Classification framework

We implemented a standardized cross-validation pipeline using gradient-boosted decision trees (XGBoost, version 1.7). A custom function performed stratified k-fold cross-validation with multiple random seeds to stabilize results across different fold partitions. For each run, samples were split into training and held-out test sets using stratified 3-fold CV, repeated across five random seeds (42, 43, 44, 45, 46). Out-of-fold predicted probabilities were averaged across seeds to obtain a robust consensus score for each sample.

XGBoost classifiers were trained with the following hyperparameters: 500 estimators, maximum tree depth of 4, learning rate of 0.05, subsampling rate of 0.8, column subsampling rate of 0.8, and L2 regularization parameter (λ) of 1.0. The logloss objective was used, and all models were trained in parallel (n_jobs = -1).

#### Performance evaluation

Performance was summarized using receiver operating characteristic (ROC) analysis. ROC curves were plotted side-by-side across the three feature spaces, and 95% confidence intervals were estimated by 1,000 bootstrap resamples at fixed false positive rate thresholds.

## ACKNOWLEDGEMENTS

This work was supported by the American Academy of Otolaryngology–Head and Neck Surgery Foundation through the Alando J. Ballantyne Research Pilot Grant. H.G. is a Core Investigator at Arc Institute, and his research is supported by Arc. We thank Chiara Ricci-Tam for her artistic illustrations. We thank Brian Plosky for his careful reading of the manuscript and for providing thoughtful and constructive feedback.

## SUPPLEMENTARY FIGURES

**Supplementary Figure 1.**
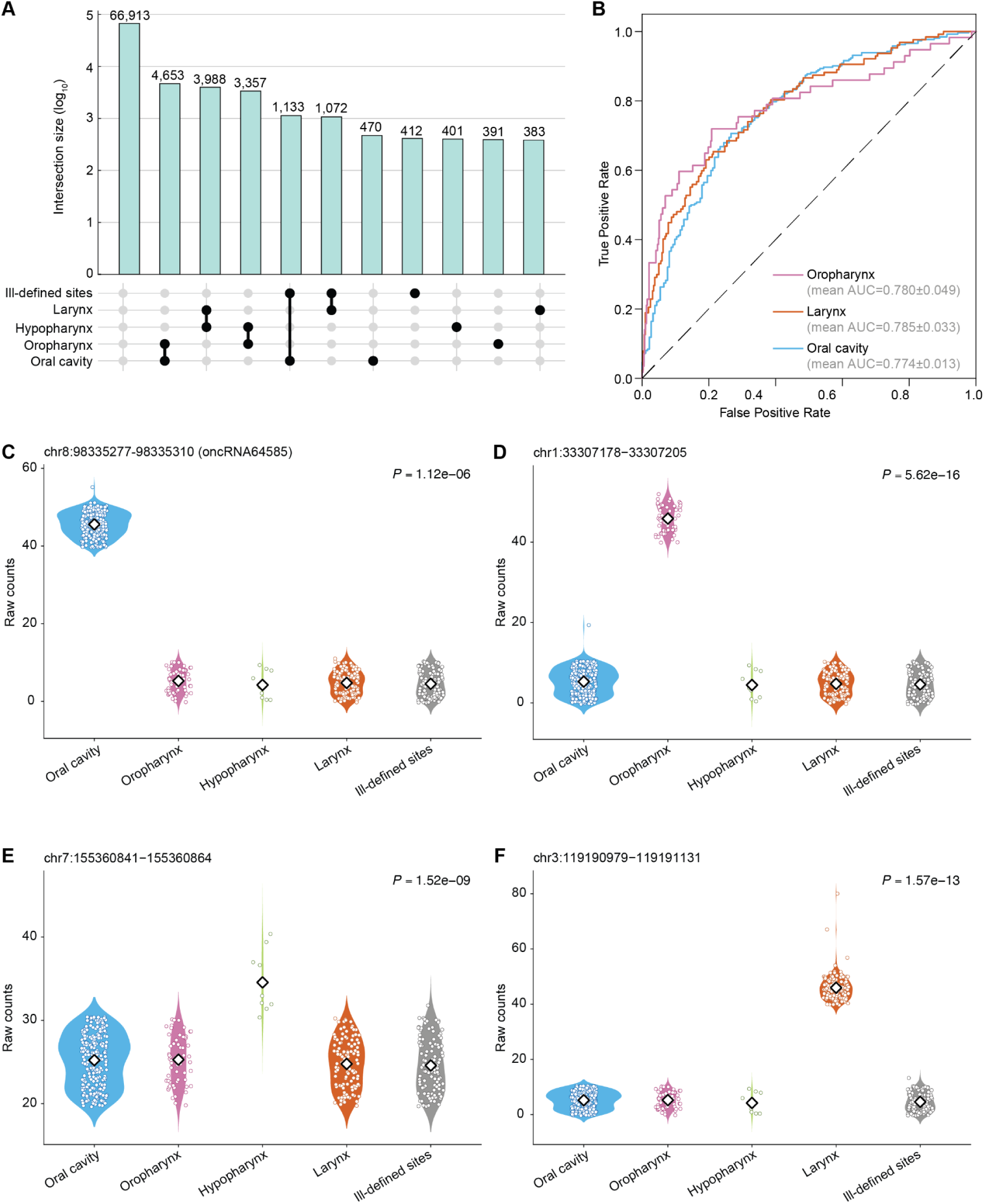
HNSCC can be stratified by their oncRNA profile. **(A)** Number of oncRNAs associated with each primary site of HNSCC, depicted as an UpSet plot. **(B)** ROC curves for XGBoost multiclass classifiers that predict the site specificity based on oncRNA presence/absence fingerprints averaged across held-out validation sets in a 3-fold cross validation setup. The vertical blue bars represent the oncRNA counts across one or more cancers with the exact numbers included at the top.**(C), (D), (E), (F)** Exemplary site-specific oncRNA loci along with their expression patterns for oral cavity, oropharynx, hypopharynx, and larynx respectively.

**Supplementary Figure 2.**
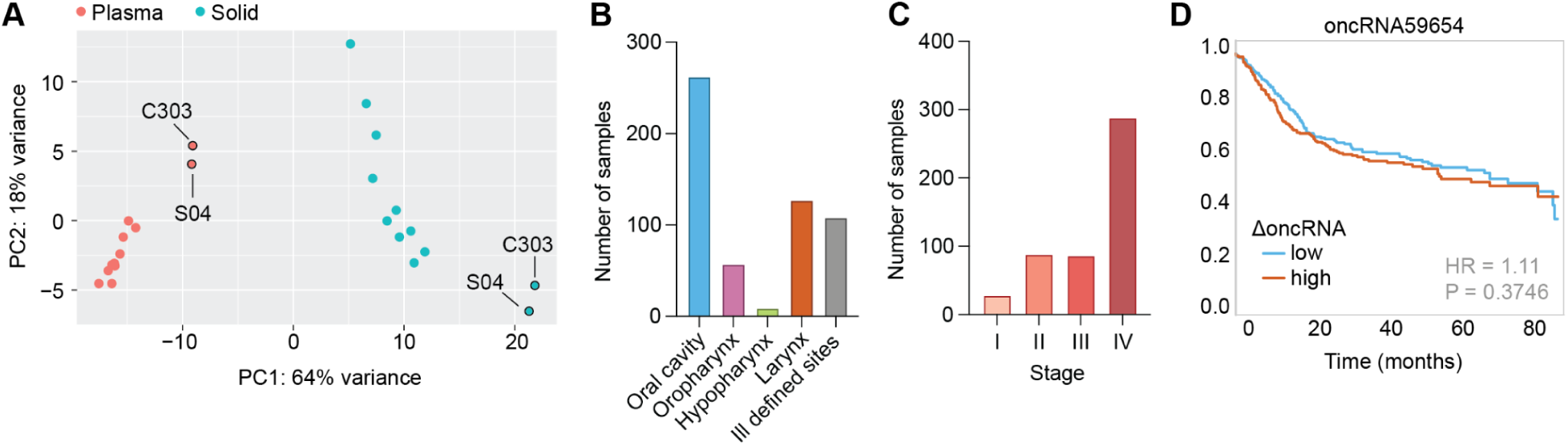
oncRNA64585 expression level corelates directly with disease progression and prognosis. **(A)** Principal component analysis (PCA) plot of smRNA-seq analysis for oncRNA-profiling in an institutional cohort of oral cavity cancer patients. C303 and S04 are non-squamous cell carcinoma cancer patients serving as negative controls. **(B)** Box plot showing distribution of patients with different primary sites of head and neck cancers.**(C)** Box plot showing distribution of patients at various stages. **(D)** Survival plot comparing patients with higher *oncRNA59654* expression (2-fold higher than the mean) versus lower expression (2-fold lower than the mean).

**Supplementary Figure 3.**
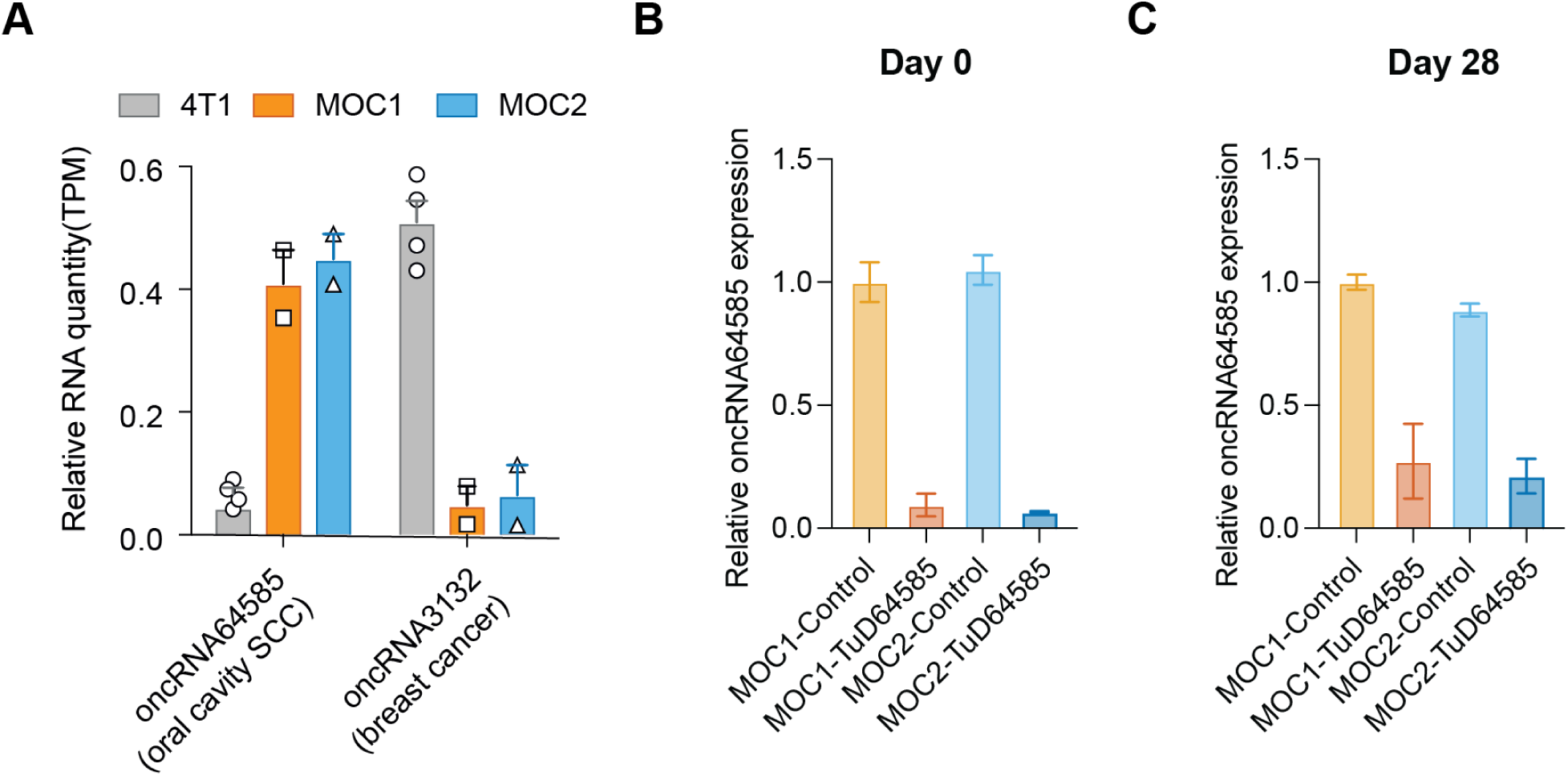
*oncRNA64585* expression promotes tumor growth. **(A)** Bar plots showing the expression levels of OCSCC-specific oncRNA64585 and BRCA-specific oncRNA3132 in 4T1 (breast cancer mouse cell line), MOC1, and MOC2. **(B)** and **(C)** Stem-loop qRT PCR levels of *oncRNA64585* prior to tumor implantation (day 0) and after explantation (day 28) respectively.

**Supplementary Figure 4.**
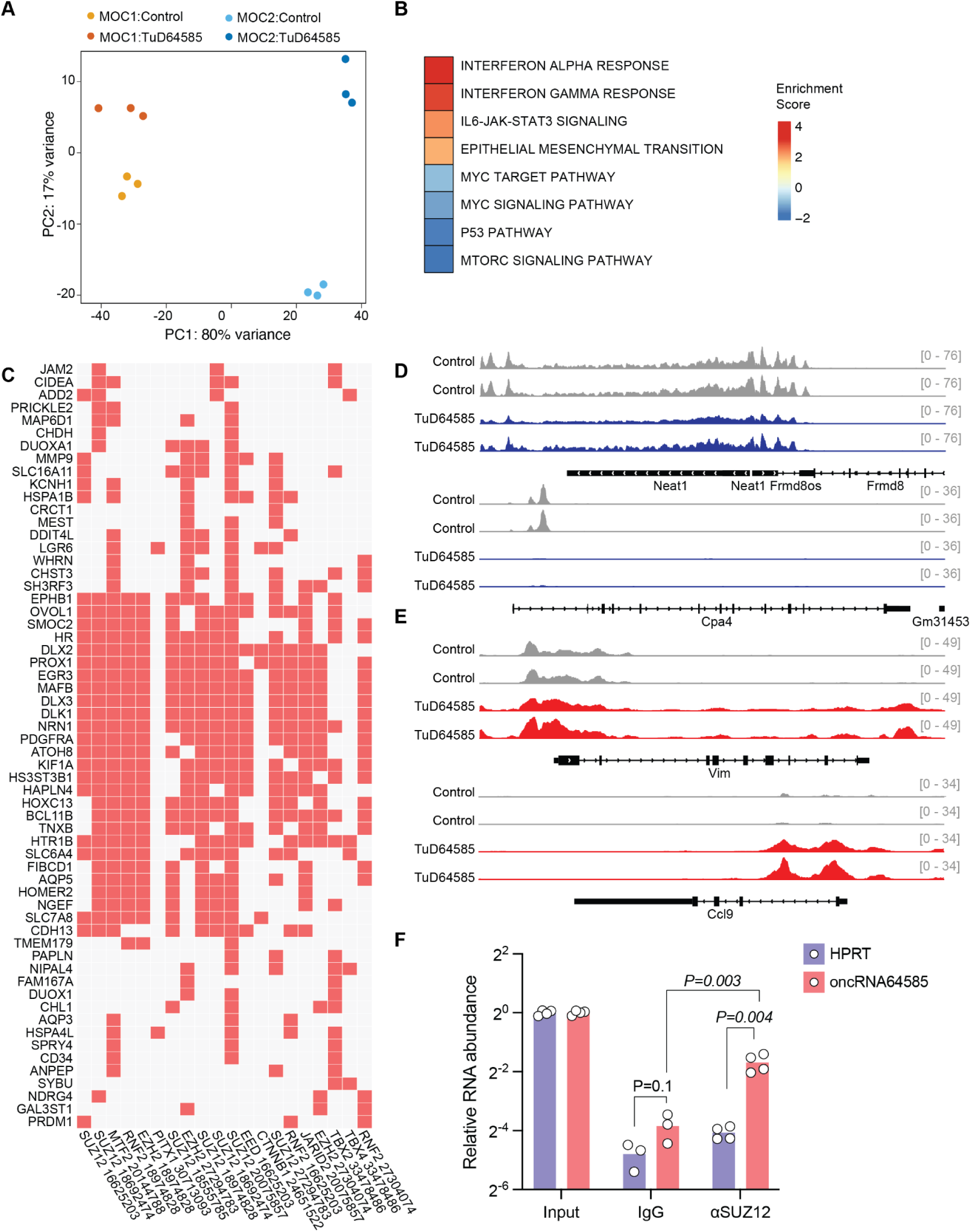
Global transcriptional and chromatin accessibility disruption as a result oncRNA64585 loss-of-function. **(A)** Principal component analysis (PCA) plot of RNA-seq analysis for MOC1 and MOC2 control or oncRNA64585 KD. **(B)** Heatmap showing enrichment of hallmark pathways of differentially expressed genes as a function of oncRNA64585 KD. **(C)** Enrichr clustergram showing enrichment of predicted co-regulators (top row) and their targets for down-regulated genes. **(D)** and **(E)** Genome-browser tracks showing two biologically replicated examples of ATAC-seq peaks near top down-regulated genes (*Neat1*, *Cpa4* in blue), or upregulated genes (*Vim*, *Ccl9* in red). **(F)** CLIP-qPCR performed using antibodies against SUZ12 (αSUZ12) or IgG control showing enrichment of *oncRNA64585* in SUZ12 immunoprecipitates compared with IgG.

**Supplementary Figure 5.**
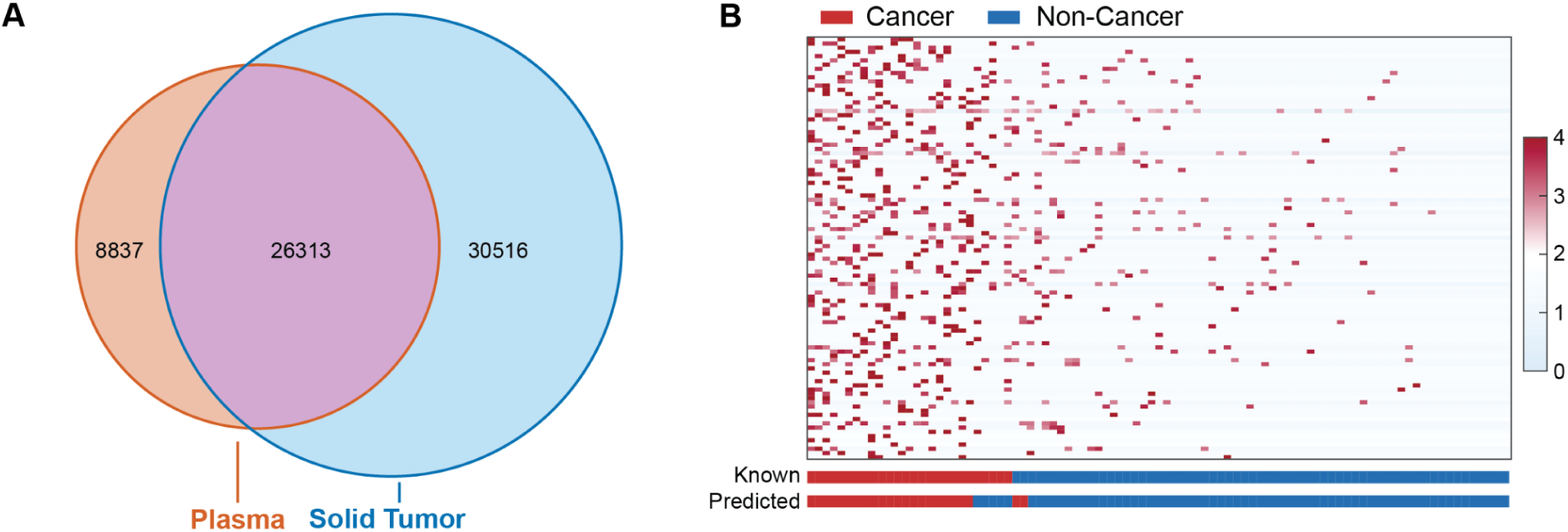
OncRNA profiles can predict cancer prior to diagnosis using liquid biopsies. **(A)** Venn diagram demonstrating shared oncRNAs between solid tumors and plasma. **(B)** Heatmap showing expression level of the top differentially expressed oncRNAs with comparison between known and predicted outcomes.

